# Use of Chitin:DNA ratio to assess growth form in fungal cells

**DOI:** 10.1101/2023.08.18.553835

**Authors:** Andrea Kovács-Simon, Helen N. Fones

## Abstract

Dimorphism, the ability to switch between a ‘yeast-like’ and a hyphal growth form, is an important feature of certain fungi, including important plant and human pathogens. The switch to hyphal growth is often associated with virulence, pathogenicity, biofilm formation and stress resistance. Thus, the ability to accurately and efficiently measure fungal growth form is key to research into these fungi, especially for discovery of potential drug targets. To date, fungal growth form has been assessed microscopically, a process that is both labour intensive and costly. Here, we unite quantification of the chitin in fungal cell walls and the DNA in nuclei to produce a methodology that allows fungal cell shape to be estimated by calculation of the ratio between cell wall quantity and number of nuclei present in a sample of fungus or infected host tissue. Using the wheat pathogen *Zymoseptoria tritici* as a test case, with confirmation in the distantly related *Fusarium oxysporum*, we demonstrate a close, linear relationship between the chitin:DNA ratio and the average polarity index (length/width) of fungal cells. We show the utility of the method for estimating growth form in infected wheat leaves, differentiating between the timing of germination in two different *Z. tritici* isolates using this ratio. We also show that the method is robust to the occurence of thick-walled chlamydospores, which show a chitin:DNA ratio that is distinct from either ‘yeast-like’ blastospores or hyphae.

## Introduction

A subset of fungi are capable of dimorphic growth. These fungi show budding growth, but under certain conditions, switch to a hyphal growth form. This dimorphic switching is often associated with temperature [1–3], but is also often seen in pathogenic fungi, in which hyphae may be produced in response to host cues and are often essential for pathogenicity or for full virulence[2, 4, 5]. Well-studied examples of dimorphic pathogenic fungi include the opportunistic human pathogen *Candida albicans*[6], the causal agent of Dutch-Elm disease *Ophiostoma novo-ulmi*[7] and the so-called ‘zombie-ant’ fungus, *Ophiocordyceps unilateralis*[8]. Hyphal growth has a number of important functions in these pathogenic fungi, including resource location, penetration and invasion of host tissues, and defence evasion[4]. Understanding the cues underlying the dimorphic switch, its timing and the extent of switching under various conditions is therefore important in understanding pathogen virulence in these fungi[1, 4, 5].

*Zymoseptoria tritici* is a plant-pathogenic fungus, the causal agent of ‘Septoria triciti Leaf Blotch’ (STB) in wheat. This common disease of temperate-grown wheat is a major source of yield losses and fungicide requirement, even in elite, resistant wheat varieties[9]. Along with many other Dothiomycete plant pathogens, this fungus is dimorphic, capable of both budding growth (blastosporulation, pycnidiosporulation) and hyphal growth. *In planta*, hyphal growth is usually initiated by the detection of a host leaf surface, and is essential for pathogenicity, as the fungus effects entry into the leaf *via* hyphal penetration of stomata[10, 11] or wounds[12, 13]. *In vitro*, hyphal growth is seen most readily in low nutrient environments such as water agar, although there is a complex interplay between both growth forms, with reversion to budding growth often seen in mature hyphae[14, 15], and this interplay also occurs *in planta*[14]. Other factors thought to play a role in determining growth form *in vitro* and *in planta* include host cues such as compounds found on the surface of a wheat leaf: sucrose, glycerol, alkanes and long-chain fatty acids[5]. There is also evidence that the switch to hyphal growth is partially regulated *via* the light sensitive white collar 1 protein[5, 16].

Growth form *in planta* is very variable between isolates of *Z. tritici*, especially during the early, epiphytic phase of plant colonisation[17, 18]. Not only does the amount, distribution and duration of epiphytic growth vary, but so too does the growth form adopted by the fungus[18, 19]. As well as the links between growth form and virulence / pathogenicity, there is also a link between growth form and biofilm formation in *Z. tritici*, which in turn is linked to stress resistance[19]. Dimorphic switching, and, more broadly, spore germination to form hyphae, are also fundamentally important processes in other fungi, including economically important plant pathogens and clinically important human pathogens[1, 2, 4]. For these reasons, determination of fungal growth form can be an important part of research tasks including forward genetic screens looking for avirulent or non-pathogenic mutants. Assessment of growth form is relatively trivial *in vitro*, but becomes more complex when considering the effect of the host environment upon fungal growth and development. For example, to assess the variation in growth form seen in field isolates of *Z. tritici* with a view to relating this to virulence or gene expression requires microscopic analysis of multiple fields of view containing multiple fungal cells at a range of timepoints during infection[12, 17, 18]. Due to the three dimension nature and complexicity of host tissues, this is best achieved using fluorescently tagged fungus and confocal microscopy[20–22]. Thus, assessment of fungal growth form *in planta* is costly and labour intensive.

Here, we propose a method for assessing fungal growth form which combines two methods for fungal quantitation from *in vitro* samples or infected host tissues to provide a ratio that reflects the shape of the average fungal cell in the sample. DNA is quantified to give a proxy measurement for the number of fungal cells in the sample, and chitin is quantified to provide a measurement of the amount of fungal cell wall present. In principle, a theoretical, perfectly round cell would contain 2*π*r units of chitin (cell wall; circumference) per unit of DNA (nucleus, one per cell). Any deviation from a perfect circle will increase the chitin measurement without changing the DNA measurement, and an elongated hypha would therefore show a chitin:DNA ratio (CDR) that would be much higher than that of any ‘yeast-like’ cell or spore (Fig. 1). We modify the method for chitin quantitation in infected leaves developed by Ayliffe *et al*.[23] to allow chitin to be measured in a range of *in vitro* and *in vivo* samples at medium-high throughput. For DNA quantitation, we use standard methods - DNA extraction followed by Qubit measurement of DNA concentration for pure fungal samples produced *in vitro*, or qPCR for specific quantitation of fungal DNA from infected host tissues. To validate the method, we compare chitin:DNA ratios to polarity indices (cell length/width) calculated from confocal scanning microscopy and image analysis conducted on samples of the same tissue. A linear relationship between CDR and polarity index for *in vitro* and *in planta* samples is demonstrated, and a suggested workflow for this methodology is provided.

**Figure 1.**
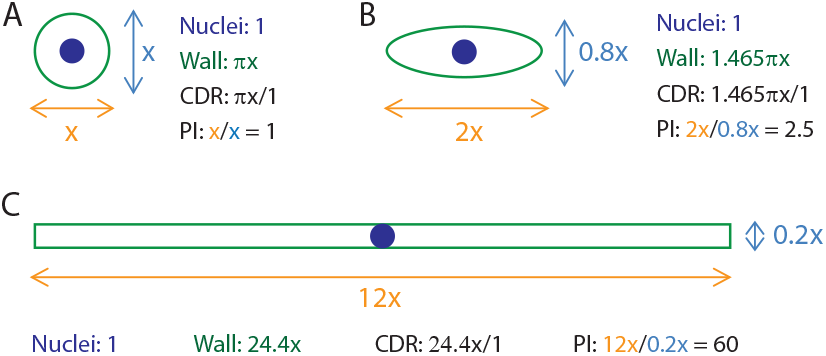
Theoretical relationship between CDR and polarity index. A theoretical perfectly round cell with a diameter x is shown (A). The cell has a polarity index of x/x = 1 and a circumference of *π*x. It has 1 nucleus. CDR is therefore *π*x/1 (*∼*3.14x). An ovoid cell, such as many fungal spores, with a length of 2x and width 0.8x (B), has a polarity index of 2x/0.8x = 2.5, a circumference of *∼*1.465*π*x and 1 nucleus. CDR is therefore *∼*1.465*π*x. Meanwhile, a hyphal cell (C), length 12x and width 0.2, has a much higher polarity index of 12/0.2 = 60 and a wall length of 2*12+0.2*2 = 24.4. Since it still has one nucleus, CDR also = 24.4. As a result, CDR is in theory a good proxy for cell shape.

## Results

### Measured Chitin:DNA ratio of *Z. tritici* cells is correlated to their polarity index

*Z. tritici* IPO323 was grown in a serial dilution of YPD liquid media. Dilution of YPD leads to an increase in hyphal growth form (Fig. 2A) (ANOVA: P < 0.0001). Cells were harvested for measurement of chitin content, DNA content and for visualisation by confocal microscopy using propidium iodide to stain cell membranes. Polarity indices were calculated and are shown plotted against measured Chitin:DNA ratios (CDRs) in Fig. 2B.

**Figure 2.**
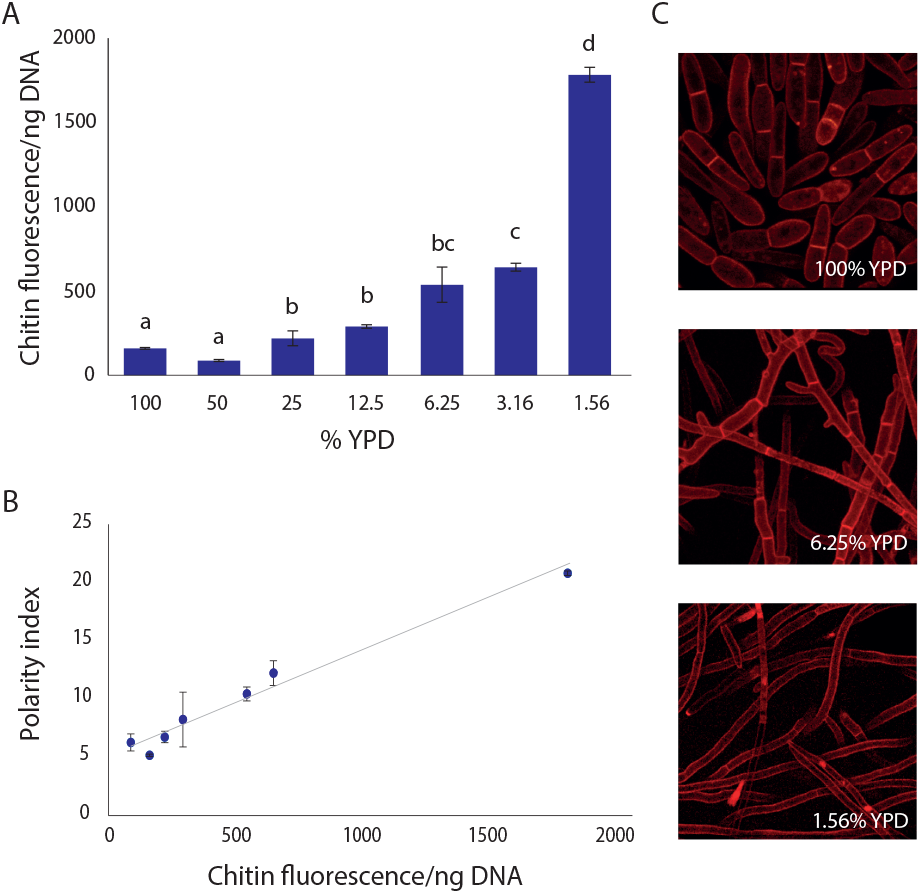
*Z. tritici* cells grown in a YPD diltuion series show closely correlated increases in both CDR and polarity index as they switch to hyphal growth at low nutrient concentrations. *Z. tritici* IPO323 cells were cultured in a 2x serial dilution of YPD broth from 100% to 1.56% for 7 days. Cultures were centrifuged and a sample of cells stained with 0.1% propidium iodide and viewed by confocal microscopy. The remaining cells were lyophilised, homogenised and divided between chitin and DNA quantitation assays from which CDR was calculated. Three technical replicates were used in each assay, and the entire experiment carried out three times. Images obtained by confocal microscopy were used to measure cell length and width and calculate polarity indices. For each media dilution, at least 5 fields of view were imaged and all cells within an image were measured. A minimum of 50 fungal structures (single or multicellular) were measured for each dilution. In multicellular structures, each cell was measured individually. CDR increased as the culture medium became more dilute (A; ANOVA, p < 0.0001), with a tight correlation existing between CDR and polarity index (B; Regression analysis, P < 0.0001, R^2^ = 0.984). Examples of the growth forms seen in the different media concentrations are shown in C; 1- or 2-cell structures predominate in 100% YPD, but the hyphal form is promoted by dilution of the media and cells become longer and thinner. Values in A and B are means of full experimental repeats (biological replicates) and error bars represent SE. Letters above bars represent significant differences in means according to Tukey’s simultaneous comparisons.

There is a tight correlation between average cell polarity indices and measured chitin:DNA ratios for each YPD concentration (R^2^ = 0.984; Regression analysis P < 0.0001). Example cells are shown in Fig. 2C. It is clear that the differences in CDR and in polarity index can be attributed to the increase in the proportion of cells in the hyphal growth form seen in low nutrient conditions. Hyphal cells have much higher polarity indices and present a much higher surface area per nucleus, giving the increased CDR that was measured. The close relationship between average polarity index and CDR indicates that CDR can be used as a proxy for polarity index, and thus for the proportion of cells in each growth form, in these samples.

To confirm the applicability of this result across multiple *Z. tritici* isolates, this assay was repeated using isolate DK09_F24. This isolate, unlike IPO323, produces hyphal growth on YPD agar within 7 days. Cultures rapidly enter a tough, wrinkled, melanised form reminiscent of the growth seen as a precursor to *in vitro* pycnidiation[24, 25]. Blastospores are produced in shaken liquid cultures. CDR and polarity index were calculated for blastospores and hyphae of this isolate and both pairs of measurements are shown in Fig. 3. Both polarity index and CDR are significantly greater for hyphal cells (*t* -tests: P = 0.0016 (polarity index); P = 0.0014 (CDR)), matching the results for IPO323.

**Figure 3.**
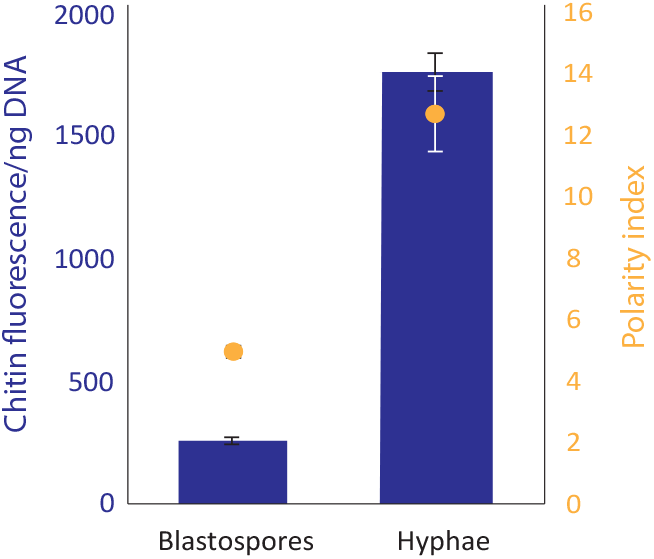
The relationship between CDR and polarity index holds when measured in an alterative *Z. tritici* isolate. *Z. tritici* isolate DK09_F24 was grown either in shaken YPD broth to produce blastospores or on YPD plates. CDR was calculated following chitin and DNA quantitation from two independent sets of three replicate samples. Polarity index was calculated following confocal microscopic visualisation of samples, from which a minimum of 50 cells were measured across a minimum of 3 randomly selected fields of view for each sample. Values are means and error bars show SE. Both CDR and polarity index are signifcantly higher in hyphal material (*t* -tests, P = 0.0014, P = 0.0016).

### Clamydospores can be detected through a distinctively high CDR

A potential complication to the use of CDR as a proxy for growth form is that *Z. tritici* has a third growth form - the chlamdyospore, described by Francisco *et al*. (2019)[14]. Chlamydospores are round and would be expected to present a low CDR based upon their shape. However, the thickness of their chitinous cell wall, reported to be four times that of blastospores[14], means that they are likely to skew the CDR of a mixed culture. To determine whether the relationship between CDR and polarity index is compromised by the presence of chlamydospores in a culture, we investigated cultures of the isolate SE13, which readily produced chlamyodospores when maintained on YPD agar. SE13 cultures were sampled across a lengthy timecourse, allowing for production and maturation of these spores, with CDR and polarity index calculated as before (see Fig. 4). Samples harvested at 8, 14, and 22 days old showed little change in CDR (*∼*100 to *∼*250; Fig. 4A) followed by a large step-change to a CDR of >3500 in 28 and 35 day old cultures. This value is over twice that seen in highly hyphal cultures in IPO323 (Fig. 2) or DK09_F24 (Fig. 3). This increase in CDR in the older cultures was significant (ANOVA with Tukey’s simultaneous comparisons; P < 0.0001). No concomitant change in polarity index was seen with this leap in CDR (Fig. 4B); in fact, across the timecourse, variation in mean polarity index was small and not explained by fungal age at harvest (slope of linear regression = 0.0003; R^2^ = 0.476; P = 0.359). Thus, for these samples, the relationship between CDR and polarity index did not hold. However, the number of chlamydospores per mm^2^ of microscopic images increased with culture age (Fig. 4C) (ANOVA with Tukey’s simultaneous comparisons; P = 0.0054) and when chlamydospores per mm^2^ was plotted against CDR (Fig. 4D), it could be clearly seen that the presence of chlamydospores explains the change in CDR where polarity index does not. Representative images of cultures at 8, 28 and 35 days old are shown in (Fig. 4E). These data together indicate that chlamydospores are distinguishable from both blastospores and hyphal cells by their much higher CDR.

**Figure 4.**
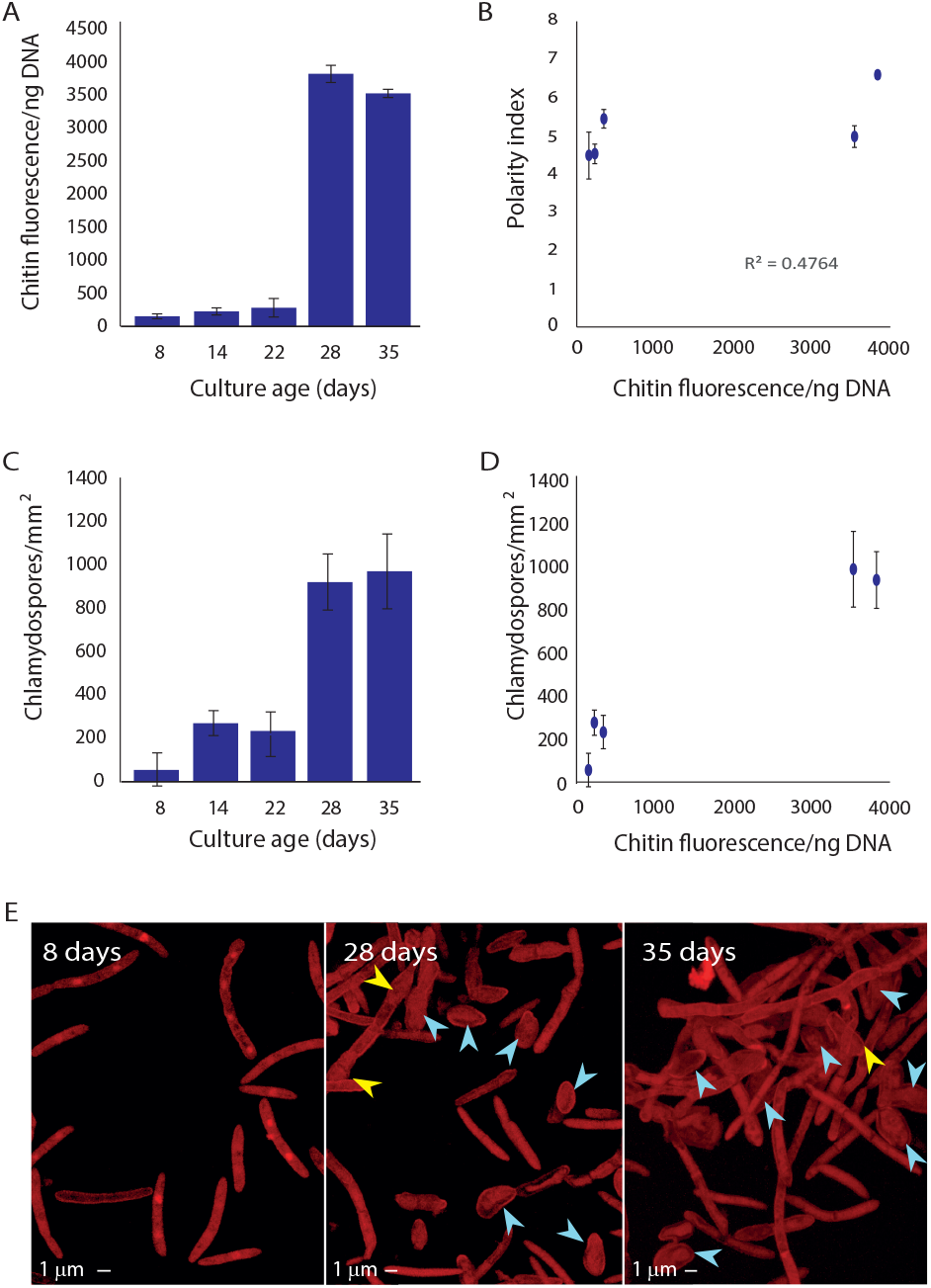
Chlamydospores affect chitin measurements and produce distinctively high CDR values. *Z. tritici* isolate SE13 was inoculated onto YPD agar plates and maintained at 20 °C for > 40 days. Chitin and DNA concentrations were measured from triplicate samples and CDR calculated at the indicated intervals (A). Samples of fungus were stained with propidium iodide and 5 randomly selected fields of view imaged at each timepoint using confocal microscopy. Polarity index was calculated from 10 randomly selected cells, or all cells, if fewer than 10, in each image (B) and chlamydospores enumerated in each image (C). There is a significant relationship between culture age and CDR (A; ANOVA, P < 0.0001), but polarity index did not vary with CDR (B; Regression, slope = 0.0003, R^2^ = 0.476; P = 0.359). However, the number of chlamydospores/mm^2^ was dependent upon culture age (C; ANOVA, P = 0.0054), explaining the relationship between culture age and CDR by the increased number of chlamydospores in older cultures (D). Representative images of 8 day old *vs*. 28 or 35 day old cultures are shown in E; yellow arrows indicate developing and blue arrows mature chlamydospores. The experiment was repeated twice independently. Values in A-D are means and error bars show SE. Different letters above bars represent significantly different means in Tukey’s simultaneous comparisons.

### CDR as a proxy for fungal growth form *in planta*

Having seen that CDR can prove and appropriate proxy for growth form *in vitro*, we investigated whether CDR could be measured and shown to have the same relationship to fungal growth form when the fungus was grown *in planta*. The wheat cultivar Galaxie was inoculated with either the *Z. tritici* isolate IPO323 or IPO94629, or strains of these isolates expressing a GFP construct expressed at the plasma membrane (SSO1GFP; [26]). Both isolates are virulent on Galaxie[18]; Chitin and DNA quantitation and calculation of CDR was carried out using leaf samples inoculated with the wildtype IPO323, while leaves inoculated with the GFP strain were harvested at the same times and viewed by confocal microscopy. As previously, polarity indices were calculated from the confocal images and plotted against CDR for the equivalent samples (Fig. 5). For both isolates, CDR increased with time after inoculation (Fig. 5A; Two way ANOVA, P < 0.0001). This increase was faster in IPO323 than IPO94629; at 3 days post inoculation (dpi), CDR for IPO94629 showed no change from day 0, while for IPO323 there was already a signficant increase (Tukey’s simultaneous comparisons, P < 0.0001). Both isolates showed a signficant increase in CDR by 12 dpi, though there was no difference between isolates at this time point (Tukey’s simultaneous comparisons, P = 0.217). As with *in vitro*-grown material, there is a close, linear relationship between CDR and polarity index for infected plant material (Fig. 5B; Regression analysis: P = 0.00024, R^2^ = 0.9747). Thus, the increase in CDR implies germination of low polarity index spores to form high polarity index hyphae, and the result at 3 dpi indicates that IPO323, but not IPO94629, has germinated by this time point. This is borne out on examination of confocal images; representative images of both isolates at 0, 3 and 12 dpi are shown in (Fig. 5C). Hyphae can be seen in both isolates at 12 dpi but only in IPO323 at 3 dpi, and at lower frequency than for 12 dpi. CDR therefore provides a good proxy for *in planta* fungal growth form.

**Figure 5.**
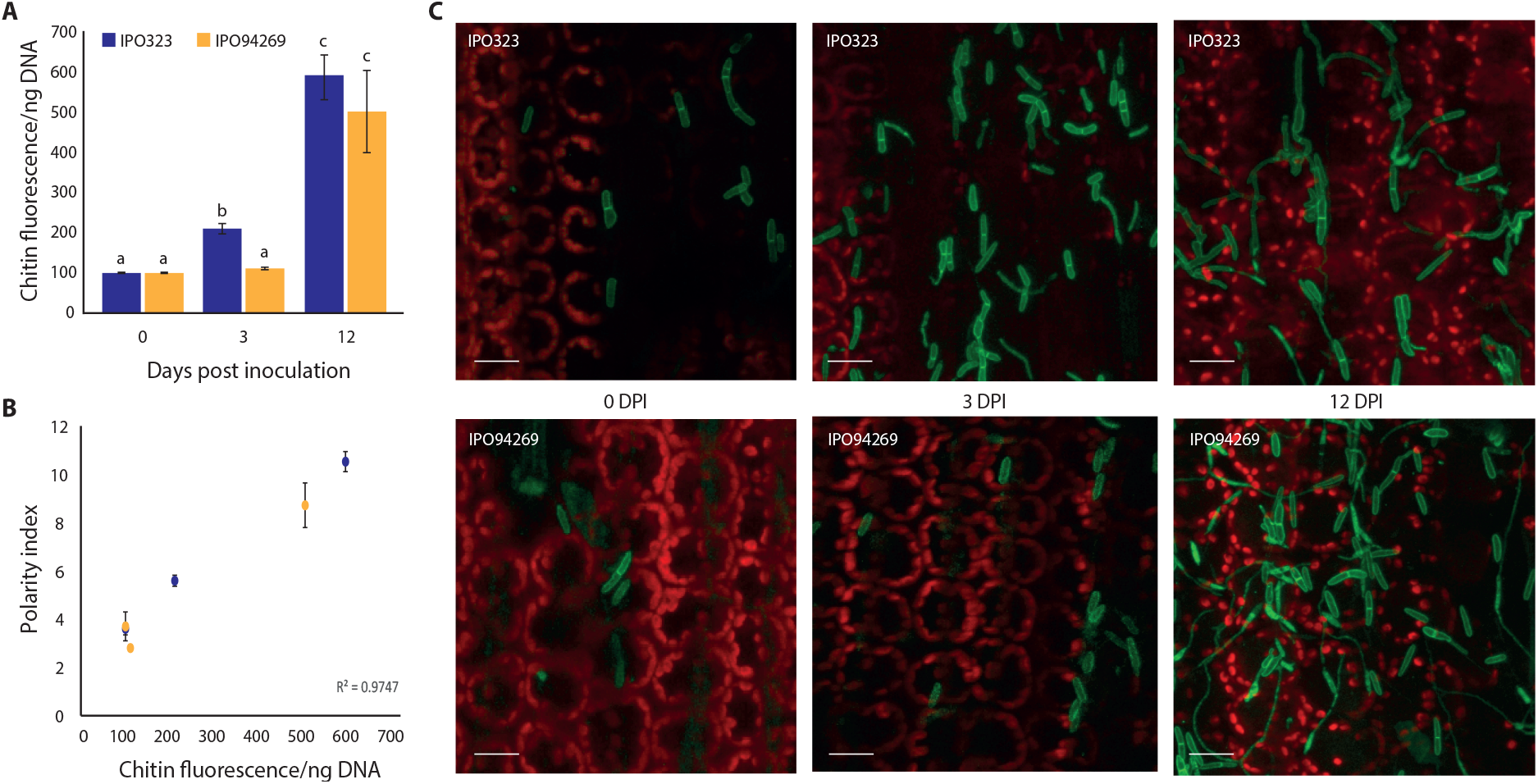
Measurement of CDR *in planta*. Wildtype and SSO1-GFP expressing strains of *Z. tritici* isolates IPO323 and IPO94629 were inoculated onto leaves of the wheat cultivar Galaxie. Samples were harvested in triplicate at 3 and 12 days post inoculation; those inoculated with wildtype isolates were used in chitin and DNA quantitation and CDR calculation (A; values are means and error bars show SE), while those expressing GFP were visualised by confocal microscopy for the calculation of polarity indices (B; IPO323, n = 121 across 3 fields of view; IPO94629, n = 117 across 3 fields of view). The experiment was repeated four times independently. Both isolates germinated on the wheat leaf, with hyphal cells visible from 3 dpi in IPO323 and at 12 dpi in IPO94629; representative images are shown in (C). CDR increased over time for both isolates (A; ANOVA, P < 0.0001 - differing letters above bars represent significant differences in Tukey’s simultaneous comparisons), but faster for IPO323, which could be seen to germinate earlier (C). As with *in vitro* samples, there was a close, linear relationship between CDR and polarity index (B; P = 0.00024, R^2^ = 0.9747).

### CDR as a proxy for growth form in other fungi

Finally, we sought to demonstrate the usefulness of CDR as a proxy for growth form beyond *Z. tritici*. We therefore measured chitin and DNA in samples of *Fusarium oxysporum f*.*sp. cubense* grown either on YPD agar to produce hyphae or in YPD broth with shaking and filtered to isolate microspores. By contrast to hyphae, microspores of *Fusarium sp*. are ovoid, with a low polarity index. As previously, we compared CDR to polarity indices calculated from confocal micrographs of propidium iodide stained cells (Fig. 6). CDR and polarity indices are shown in Fig. 6A, while examples of microspores and hyphae are shown in Fig. 6B and C, respectively. Both the increased polarity index and the increased CDR in hyphal material are highly significant (*t* -tests, P < 0.0001 (polarity index); P < 0.0001 (CDR)). All chitin and DNA quantitation was carried out in triplicate and the experiment repeated four times independently. Polarity indices were calculated from 3 randomly selected fields of view from which a total of 50 (spores) or 30 (hyphae) cells were analysed.

**Figure 6.**
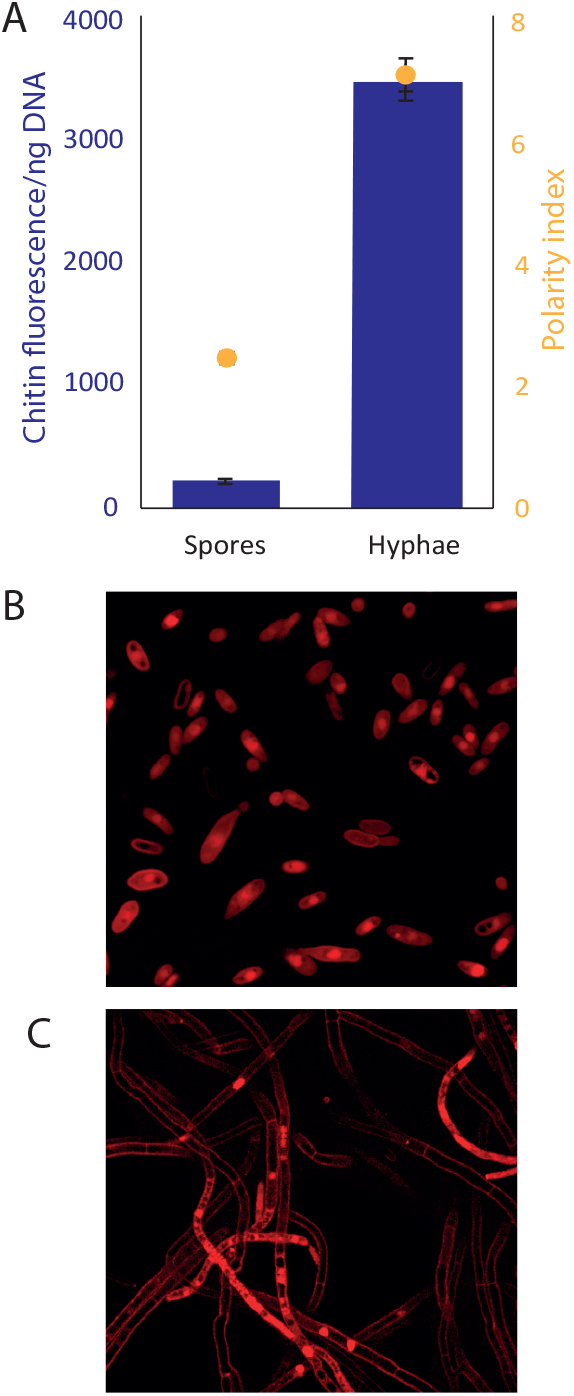
*F. oxysporum f*.*sp. cubense* shows the same relationship between CDR and polarity index as *Z. tritici*. *F. oxysporum f*.*sp. cubense* was grown either on YPD agar or in YPD broth with 200 rpm shaking. Fungal mycelium was scraped from plates, suspended in YPD broth and vortexed vigorously to loosen any spores. The mycelium was then filtered through 2 layers of miracloth and the fitrate discarded, leaving mycelium. Liquid cultures were similarly filtered but the microspore-rich filtrate was retained and the mycelium discarded. Chitin and DNA in these samples were measured and aliquots stained with propidium iodide, visualised by confocal microscopy and polarity indices calculated. Values are means and error bars show SE. Both CDR and polarity index are signifcantly higher in hyphal material (*t* -tests, P = 0.0014, P = 0.0016).

## Discussion

In this work, we have described a methodology for estimating fungal growth form based upon simple assays to quantify DNA and chitin. We exploit the difference in the ratio of length of cell periphery to number of nuclei that occurs when cells change shape, hypothesising that this must translate into differences in the ratio between the amount of the cell wall constituent, chitin, and DNA (Chitin:DNA ratio, CDR) in samples of fungus. Using the wheat pathogen *Zymoseptoria tritici* as our test case, we have shown that, indeed, cell polarity index in this species shows a close, linear relationship to CDR in most contexts. Polarity index, calculated according to cell length/width, is a measure of a cell’s shape where a circular cell has a polarity index of 1; the longer and thinner a cell is, the higher the polarity index.

This relationship between CDR and polarity index means that the shape of *Z. tritici* cells can be inferred from CDR alone; thus, CDR is a good proxy for *Z. tritici* growth form. Growth form is of large importance in

*Z. tritici* biology, with different functions carried out by cells in each form. The fungus is generally classed as dimorphic[11, 27, 28], although there is variability in polarity index of cells *in planta*, such that a bimodal distribution of polarity indices is more accurate. It produces spores - both sexual and asexual - composed of cells with low polarity indices[11], which germinate on the wheat leaf surface to produce long, thin hyphae[11, 28, 29]. Growth form on the leaf is fluid, with budding (variously called blastosporulation[14], micropycnidiation[11] and microcycle conidiation[12]), germination and anastomosis all occurring from both pycnidia and blastospores[14], and there is significant variation in the proportion of each seen in different isolates at different times during infection[17, 18, 30].

Measuring growth form provides information about fungal development and cell function. In *Z. tritici*, hyphae are essential for virulence[31–33] as they are responsible for stomatal penetration to gain access to the interior of the leaf, and for the colonisation of the apoplast. Conversely, epiphytic proliferation of blastospores is associated with the largely avirulent ‘NIRP’ phenotype[18], and both hyphae and blastospores play a role in biofilm formation[19]. A large research effort has been applied to understanding the environmental and genetic mechanisms that lead to the formation of hyphae, as this is a key step in host invasion and thus virulence[5, 24, 31, 32]. This is true not only in *Z. tritici*, but it other dimorphic fungal pathogens, including plant pathogens such as corn smut (*Ustilago maydis*)[34] and Dutch elm disease (*Ophiostoma novo-ulmis*)[7] as well as many important yeast and fungal pathogens of humans. In dimorphic human pathogens, as in plant pathogens, growth form is closely related to functions including host invasion, biofilm formation and virulence[3, 6, 35]. Dimorphism is therefore closely associated with virulence. Further, it is linked to stress resistance. In many species, the switch to hyphal growth is dependent on environmental factors such as temperature and nutrient availability, and different growth forms show differential abilities to withstand these stressors[19].

Given the importance of fungal dimorphism, it is common to carry out medium- or high-throughput screens for fungal growth form, for instance to identify mutations that switch to hyphal growth more or less readily and may be compromised in host invasion or stress resistance. Such screens are often used to identify genes that may be appropriate anti-fungal targets [33, 36, 37]. However, *in vitro* screens for fungal growth form do not always identify strains that are most affected in the host environment. In the case of *Z. tritici*, the host factors that promote germination to hyphae are not fully elucidated [5, 30]. One of the key triggers known to induce hyphal growth *in vitro* - cultivation temperatures of above 28 °C[5, 14, 36] - is unlikely to be present in temperate grown wheat during the Spring months, when *Z. tritici* infections are established[38–40]. Such considerations make the *in planta* screening of *Z. tritici* growth form desirable.

To date, *in planta* measurement of fungal growth form has relied on a combination of staining and microscopy to distinguish fungus from host tissues and identify the positions of cell walls and septae to allow the measurement of polarity index. For complex, 3D tissues, as well as to study invasive fungal hyphae, the combination of fluorescent protein expression in the fungus and confocal laser scanning microscopy has been widely used. These methods present drawbacks: they are costly, labour intensive, and therefore limited to small areas of infected host tissue and relatively low numbers of fungal cells. Measurement of polarity index from images requires complex image analysis or is, again, labour intensive. Further, not all laboratories have capacity to work with genetically modified (GFP-expressing) pathogens or access to confocal microscopy facilities.

The use of CDR offers an alternative methodology by which fungal growth form can be estimated from both *in vitro* fungal samples and infected leaf material. We have shown that the correlation between CDR and polarity index is not specific to *Z. tritici*, but holds when tested in *Fusarium oxysporum*, another Ascomycete fungus, but one which belongs to a different Class to *Z. tritici* and produces multiple spore types but no ‘yeast-like’ growth. It is therefore likely that the CDR method for estimation of fungal growth form would be applicable to all fungi and any non-chitinous host material sampled. A workflow diagram describing the CDR measurement and calculation protocol is provided in Fig. 7. A number of caveats must be considered when applying the CDR method for estimation of polarity index and thus growth form:

**Figure 7.**
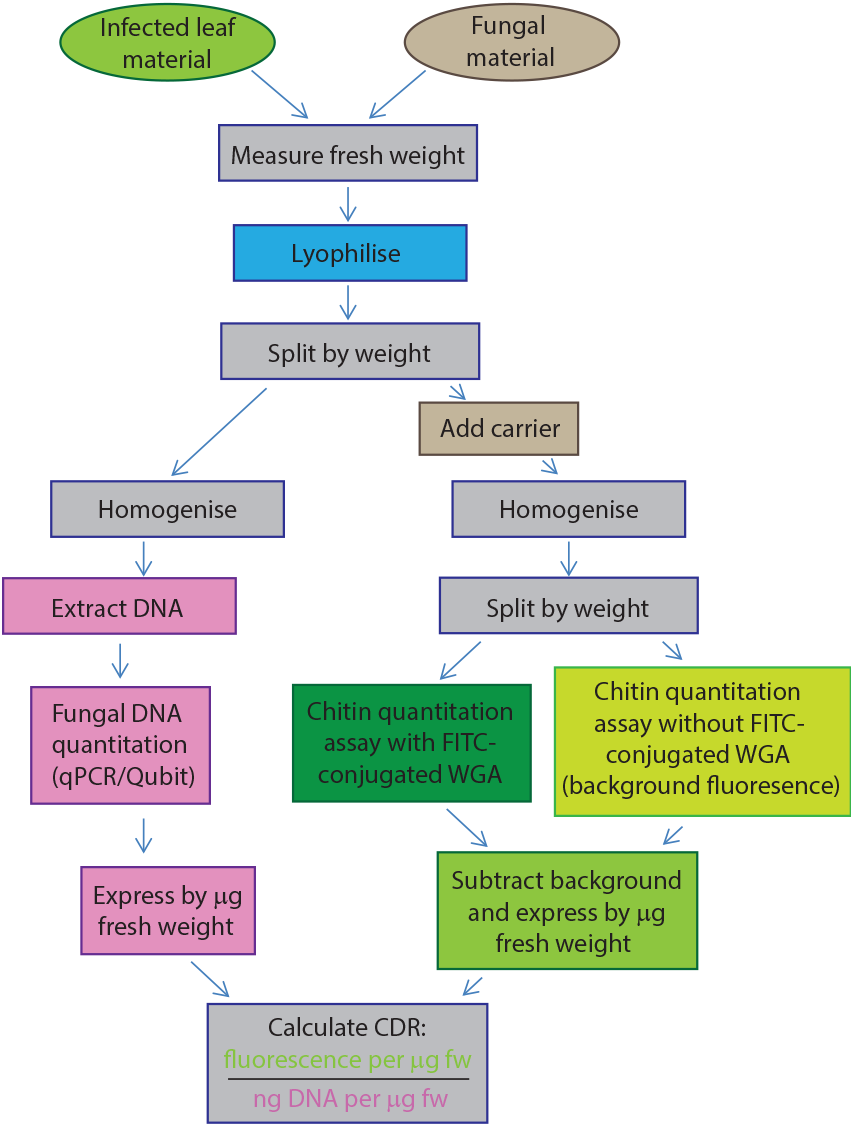
Suggested workflow for CDR calculation. Notes: 1. ‘carrier’ required only for use with small-volume samples of pure fungal material, to improve homogenisation; 2. see methods for details of chitin quantitation method, including refinements made in this work.

- Are the growth forms of the fungus of interest readily distinguished by polarity index? As shown here, polarity index and thus CDR can distinguish between hyphal and ‘yeast-like’ (blastospore) growth, or between hyphae and spores. It may not be suitable for distinguishing between multiple spore types of the same fungus; we did not attempt to distinguish, for example, micro- and macrospores of *Fusarium oxysporum*, and would not necessarily expect this to be possible, as macrospores are made up of cells with similar polarity indices to microspores. Similarly, ascospores, pycnidiospores and blastospores of *Z. tritici* are likely to be indistinguishable by CDR.
- Are there any additional growth forms that might arise, complicating the interpretation of CDR? We showed that the chlamydospores of *Z. tritici* can be detected *via* the CDR method because they produce a much higher CDR even than pure hyphal material (*∼*3500 for chlamydospores *vs ∼*1800 for hyphae). However, if applying the method to other fungi, it would be advisable to check the ranges of CDR values associated with samples of each possible growth form to ensure these do not overlap.
- Are variations in the number of nuclei per cell likely? In this work, we tested only Ascomycete fungi. Therefore, we did not consider coenocytic hyphae nor dikaryons, which may be present in other fungal phyla. The CDR method is likely to remain effective for aseptate hyphae, unless the distance between nuclei is variable.
- Is there background fluorescence in the host tissues? Since plant cell walls and chloroplasts are autofluorescent, we included unstained control samples in our experimental design. These allowed us to subtract the background fluorescence of the host tissue from each sample, in a sample-specific manner; we recommend this control. Not only are plants autofluorescent, but the degree of fluorescence can change during the activation of plant defences[41]. In line with this, we found that chlorotic leaves showed higher background fluorescence in our assays (Fig. S3).
- Fungal fluorescence? GFP-tagged fungi may not be suitable for quantifying chitin *via* FITC, as these molecules have overlapping fluorescence emission spectra. Consider an alternative fluorophore conjugate for WGA for such assays.

We recommend that the relationship between CDR and polarity index is confirmed microscopically for any new fungal species.

In summary, we have shown that the ratio of chitin and DNA in fungal cells can be used as a proxy for polarity index and thus growth form. Using the fluorescence based chitin quantitation methods developed by Ayliffe *et al*.[23] combined with specific DNA quantitation by qPCR, it is possible to estimate the quantity and growth form of fungi in infected tissues, without the requirement for extensive, labour intensive microscopic analysis. This method can be used to facilitate screens for growth form in host tissues.

## Methods

### Fungal isolates and growth conditions

*Z. tritici* isolates IPO323 and IPO94269 were kindly provided by Prof Gert Kema and Prof Gero Steinberg. IPO323 and IPO94269 were isolated in the Netherlands and have previously used in multiple studies of *Z. tritici*, with IPO323 commonly regarded as the reference isolate for this fungus[42–44]. Dk09_F24 was isolated in Denmark and kindly provided by Prof Eva Stukenbrock[45]. SE13 was isolated by Dr Helen Fones in Southern England. Strains of each isolate expressing cytosolic GFP or SSO1-GFP[26] were produced by integrating the GFP construct into the *sdi1* locus, according to the methods of Kilaru *et al*. (2015)[21], using vector plasmids kindly provided by Dr Sreedhar Kilaru. All isolates were grown on yeast-peptone-dextrose (YPD) agar at 20 °C and or liquid YPD broth at 20 °C with 200 rpm shaking. *Fusarium oxysporum f*.*sp. cubense* was kindly provided by Prof Daniel Bebber and cultured on YPD agar at 18 °C or in liquid YPD broth at 18 °C with 200 rpm shaking. Spores and hyphal material of *Fusarium oxysporum f*.*sp. cubense* and *Z. tritici* Dk09_-F24 were separated by filtering broth cultures or fungal material scraped from plates and resuspended in sterile YPD broth through 2 layers of miracloth.

### Wheat growth conditions and inoculations with *Z. tritici*

Wheat (*Triticum aestivum* cv. Galaxie) was grown on John Innes No.2 compost in a growth chamber with 12 h light, 20 °C day time temperature and 18 °C night-time temperature and 80% relative humidity. Inoculations were carried out on 14 day-old plants. *Z. tritici* blastospores were suspensed in sterile distilled water supplemented with 0.1% Tween-20, filtered through two layers of Miracloth and adjusted to 10^6^ cfu/ml. Fully expanded wheat leaves were marked to show inoculation sites and marked portions coated with spores suspensions using a paintbrush. Once inoculated, plants were kept at 100% RH for the first 72 h. Following this, plants were maintained as before.

### Chitin quantitation

Chitin quantitation followed the method presented by Ayliffe *et al*.[23], with modifications. Around 50-100 mg (fresh weight) of infected leaf tissue or 5-10 mg of pure fungal tissue per sample was harvested into 2 ml tubes, frozen in liquid nitrogen and lyophilised (Scanvac CoolSafe Touch 110-4 Freeze Dryer). The lyophilised mass of each sample was recorded and samples were stored at -80 °C until required. Lyophilised material was homogenised using a TissueLyser™ at 30 rpm for 3 mins. 600 *μ*l 1 M potassium hydroxide was added to each sample and mixed by vortexing. Homogenate suspensions were then heated to 100 °C for 30 mins. After this heat treatment, 1.2 ml 50 mM Tris was added to each tube. Samples were then centrifuged for 30 mins at 13,000 rpm and the supernatant removed. Pellets were resuspended in 800 *μ*l 50 mM Tris and allowed to equilibrate overnight 4 textsuperscriptoC. Samples were then divided into two equal volumes and 20 *μ*l of 1 mg/ml (w/v) wheatgerm agglutinin FITC conjugate (WGA-FITC) was added to one of each pair and incubated at room temperature for 15 mins. Following this incubation, 1.4 ml 50 mM Tris was added to all samples. Samples were then centrifuged for 15 mins at 13,000 rpm and pellets washed three times with 1.8 ml Tris. Washed pellets were resuspended in 400 *μ*l Tris. Triplicate 100 *μ*l samples were transferred to wells of a black 96 well plate. In addition Tris-only controls were included, in triplicate, as well as triplicate samples of Tris supplemented with 10 *μ*l of 1 mg/ml (w/v) WGA-FITC to provide a standard for comparison of fluorescence readings between plates. Fluorescence was measured in a Clariostar plate reader with 485 nm adsorption and 535 nm emission wavelengths. Chitin fluorescence was then expressed per mg (fw) of starting tissue.

### Refinements to chitin quantitation method

In order to further ensure the applicability of the method presented here in the high-throughput manner to the widest possible range of fungi and host tissues, a number of experiments designed to refine the method presented by Ayliffe *et al*.[23] were carried out.

#### Heating method

Firstly, to remove the need for an autoclave, used in the original method[23], alternative methods of heating the test samples were tested. Many labs have only one or two autoclaves which may be in high demand. Further, cycle time can be in excess of 3 h for some models, if run full. The use of an autoclave therefore presented as a throughput-limiting step in the protocol. The effect upon sensitivity of chitin detection of replacing autoclave treatment with heating for 30 mins in a heat-block at 100 °C was tested (Suppl. Fig. S1). A greater range of fluorescence readings were obtained across a dilution series of an inoculated wheat leaf sample with the heat-block method, in particular with the undiluted sample producing higher fluorescence; see Supplementary Figure S1. These results indicated that availability of chitin in the sample to the dye was better in the heat-block protocol. The heat-block protocol was thus adopted in all experiments.

#### Optimal homogenisation of fungal material

Direct quantitation of chitin from fungal samples was carried out according to the same method as for infected plant material with certain modifications. Smaller quantities of starting material are appropriate for pure fungal samples; however, small volumes of material presented difficulties during homogenisation. Initial experimentation showed variation in results per mg of fungal material much greater than that seen per mg of infected leaf material. The homogenisation step showed variable success, and samples visually assessed as poorly homogenised yielded lower apparent chitin quantities in comparison to those seen in samples visually assessed as successfully homogenised by the same protocol. Homogenisation was improved by the introduction of the lyophilisation step during sample preparation (see above). However, homogenisation success still appeared to vary with fungal growth form. Since this variation was not seen in the results from infected plant material, we hypothesised that the presence of the additional bulk of plant leaf material in these samples may have improved homogenisation efficacy, for example by preventing the aggregation of fungal material in the lid or in the tip of the tube during processing in the TissueLyser™. The inclusion of 40 mg of non-inoculated, frozen wheat leaf material prior to homogenisation of fungal material, to act as a ‘carrier’ for fungal homogenisation, was therefore tested. This greatly reduced the intra- and inter-sample variability in the fluorescence results. The inclusion of a carrier for homogenisation of fungal samples was therefore adopted. Since production of wheat leaves for this purpose is relatively labour-intensive and would not be appropriate in all settings, as well as providing a degree of background fluorescence, cellulose powder (Sigma, UK) was tested as an alternative carrier. 40 mg cellulose was added at the same point prior to homogenisation. As with the use of the leaf carrier, cellulose carrier reduced intra- and inter-sample variability; the sensitivity of the assay was unaffected; see Supplementary Figure S2. We recommend the use of cellulose as a carrier for small volume samples.

#### Accounting for background leaf fluorescence

As described above, each sample was split into two prior to the addition of WGA-FITC to one of each sample pair thus produced. Unstained controls were included, in triplicate, alongside each sample. This allowed measurement of background fluorescence coming from leaf tissue in each sample. Leaves with visually different levels of chlorosis were also assayed in order to determine whether leaf health or colour affected this background (Suppl. Fig. S3). As results differed between samples, with chlorotic leaves showing significantly higher background fluorescence at 535 nm, we concluded that the inclusion of no-stain controls is important, especially if comparing leaves infected with isolates of differing or unknown virulence, where differences in symptoms including chlorosis may be expected.

## DNA quantitation

To extract DNA, leaf tissue or fungal material was flash frozen in liquid nitrogen and homogenised in a TissueLyser™ at 300 rpm for 3 mins. DNA was extracted using a Plant/Fungi DNA Isolation Kit (Norgen) according to the manufacturer’s instructions. DNA was eluted in 100 *μ*l elution buffer and aliquots diluted 10x for use in qPCR. DNA concentrations were measured on a Qubit 4 fluorometer (Invitrogen, UK) for samples containing only fungal DNA, or by qPCR for mixed plant/fungal DNA samples, in order to specifically quantify fungal DNA. Neat and diluted DNA was stored at ^-^20 °C until required. qPCR was carried out in a QuantStudio™7 Flex Real-Time PCR System with the primer pair ST-rRNA F/R, developed for *Z. tritici* qPCR by Guo et al (2006)[46] with the cycling conditions specified in that work. A standard curve of serially diluted *Z. tritici* DNA at known concentrations was included to allow calibration of results. No template controls and no primer controls were included to confirm the absence of non-specific amplification.

### Calculation of CDR

Single samples of homogenised leaf or fungal tissue were split into equal parts by mass. Chitin fluorescence and ng DNA were then measured in these identical sample aliquots and results adjusted according to original sample mass. Comparable, per mg values were thus obtained for each measurement and the Chitin-DNA ratio (CDR) calculated simply as fluorescence per mg of sample / ng of DNA per mg of sample.

### Confocal microscopy

Fungal samples were suspended in 0.1% (v/v) phosphate buffered saline (PBS, pH 7) and stained by supplementation of PBS with 0.05% (w/v) propidium iodide (PI) to a final concentration of 0.05% (w/v). Samples of leaves inoculated with GFP-tagged strains of all fungal isolates were mounted in 0.1% (v/v) PBS. Confocal microscopy was carried out using argon laser emission at 500 nm with detection in 600–630 nm (chlorophyll/PI, red) and 510–530 nm (GFP, green), using a Leica SP8 confocal microscope with 40x oil immersion objective.

### Calculation of polarity index

Confocal micrographs showing PI-stained or SSO1-GFP expressing cells were used to identify the location of the plasma membranes and septae. The length and width of each individual cell was then measured manually in Image J[47] and polarity index calculated as length/width.

### Statistical analyses, randomisation and blinding

Statistical analyses were carried out in ‘R’[48] or excel. Details are given in figure legends, but all analyses were either *t* -tests, ANOVAs or regression analyses. Data were checked for conformation with relevant assumptions using the Shapiro-Wilk test for normality and Levene test for homogeneity of variance, as well as visual inspection of a Q-Q plot of residuals and residuals *vs*. fits plot, as appropriate. For microscopic analysis, random fields of view were selected by focusing in the top left corner of the sample followed by a random number generator based walk right and down through the sample. Only fields of view containing no fungi were rejected. During microscopic analyses the human operator was also blinded as to the polarity index that might be expected in a sample as the labels on samples were coded by a different investigator, and only related to their correct sample names after data collection.

## Supplementary Information

**Figure S1.**
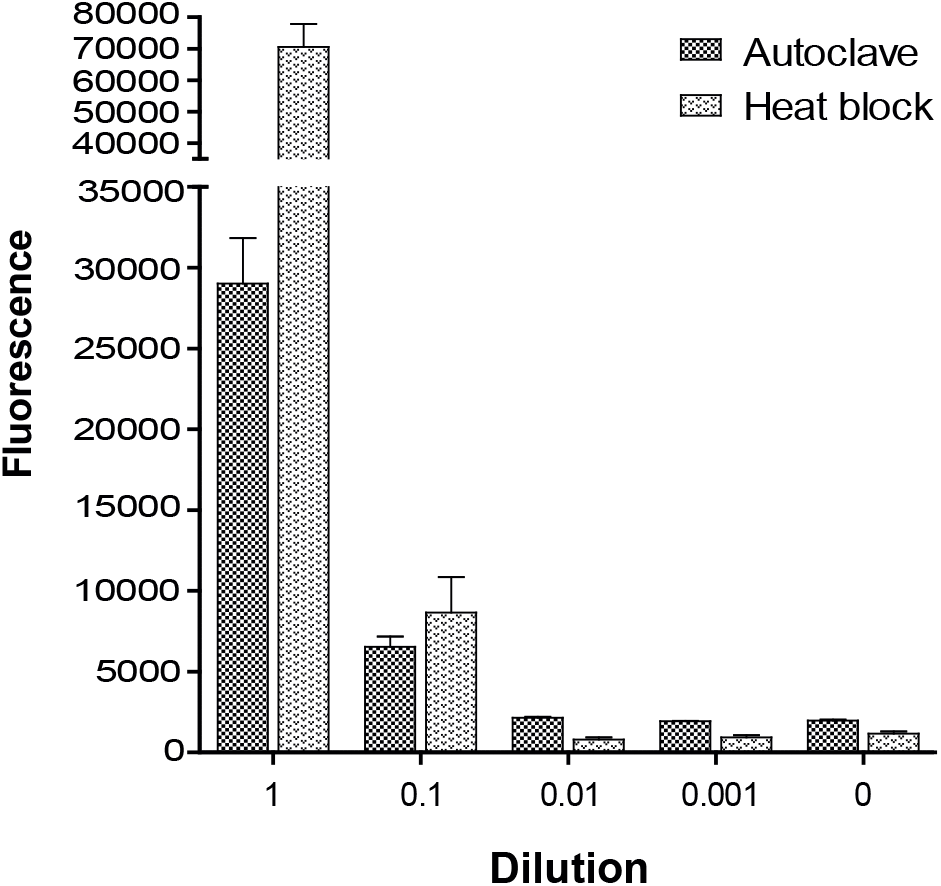
Autoclave vs heat block. 1 M KOH was added to lyophilised, homogenised leaf material and samples then either autoclaved, according to the protocol in Ayliffe *et al*[1], or heated to 100 oC in a heatblock for 30 min. A dilution series was created from the samples and the rest of the protocol followed as usual. Undiluted samples give significantly higher fluorescence when autoclaved, but other dilutions showed no differences, suggesting that the limits of detection are similar for both methods. Values are means of 4 replicates and error bars show SE.

**Figure S2.**
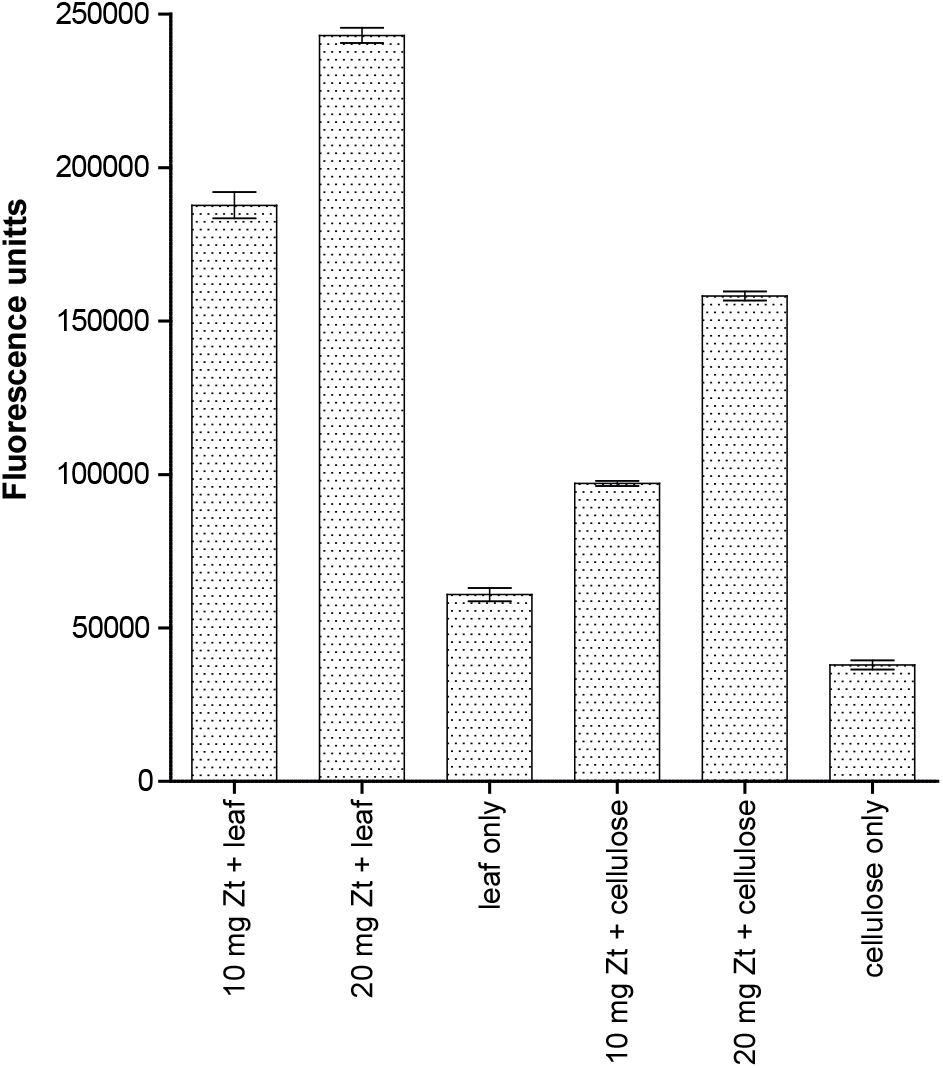
Leaf vs cellulose as carrier. We tested the use of both uninfected leaf material and purified cellulose as ‘carriers’ for small samples (10 or 20 mg) of *in vitro* grown fungus during the homogenisation step. Both sets of results show the expected increase in fluorescence when more fungus is present in the original sample. Values are means of 4 replicates and error bars show SE.

**Figure S3.**
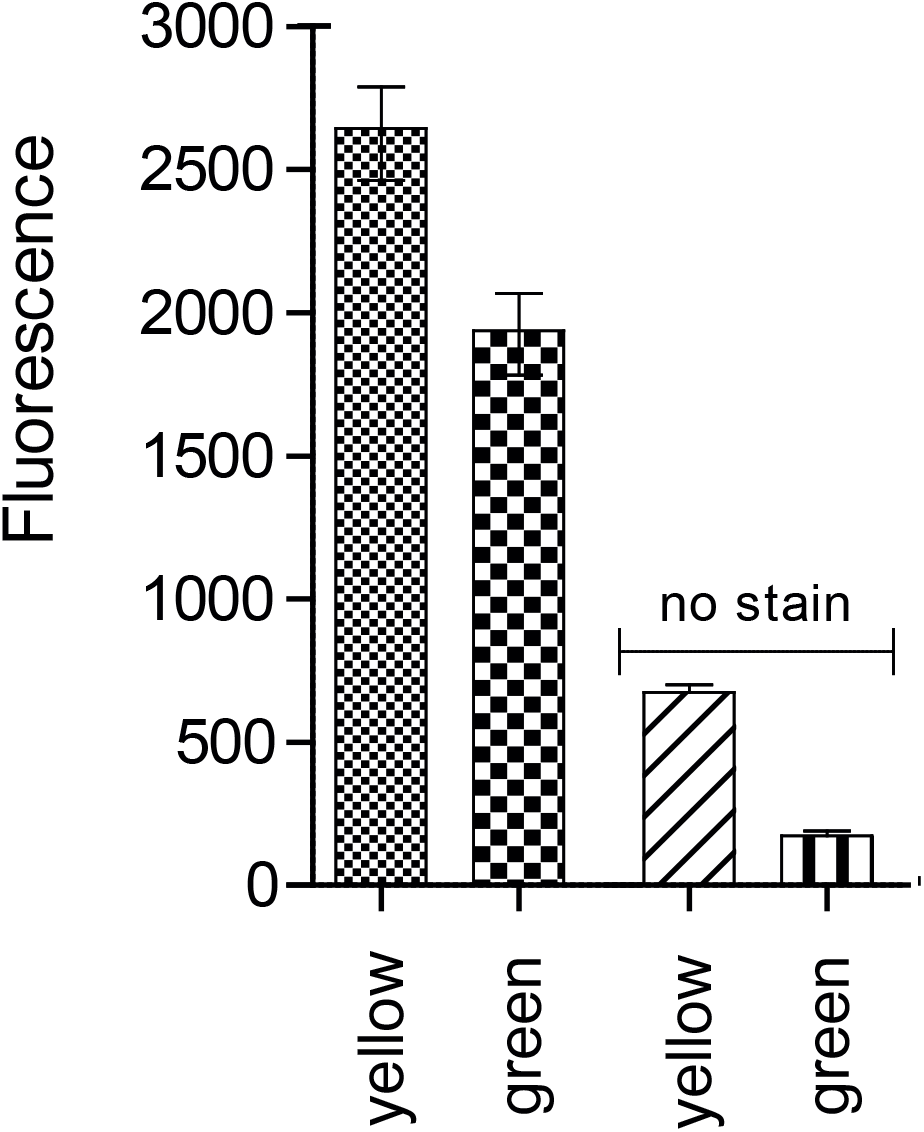
Effect of leaf health on background fluorescence. We carried out chitin quantitation assays on samples of inoculated leaf material immediately after inoculation. For inoculation we selected leaves which could easily be visually distinguished on the basis of colour. Combined with the inclusion of the ‘no dye’ controls for each sample, this allowed us to determine whether apparently healthy green leaves provided a different level of background fluorescence than senescing, yellow leaves. Fluorescence values from yellow leaves were consistently higher. Values are means of 4 replicates and error bars show SE.

